# Disrupting folate metabolism alters the capacity of bacteria in exponential growth to develop persisters to antibiotics

**DOI:** 10.1101/335505

**Authors:** Jasmine Morgan, Matthew Smith, Mark T. Mc Auley, J. Enrique Salcedo-Sora

## Abstract

Bacteria can survive high doses of antibiotics through stochastic phenotypic diversification.We present initial evidence that folate metabolism could be involved with the formation of persisters. The aberrant expression of the folate enzyme gene *fau* seems to reduce the incidence of persisters to antibiotics. Folate impaired bacteria had a lower generation rate for persisters to both antibiotics ampicillin and ofloxacin. Persister bacteria were detectable from the outset of the exponential growth phase in the complex media. Gene expression analyses showed tentatively distinctive profiles in exponential growth at times when bacteria persisters were observed. Levels of persisters were assessed in bacteria with altered, genetically and pharmacologically, folate metabolism. This work shows that by disrupting folate biosynthesis and usage, bacterial tolerance to antibiotics seems to be diminished. Based on these findings there is a possibility that bacteriostatic antibiotics such as antifolates could have a role to play in clinical settings where the incidence of antibiotic persisters seem to drive recalcitrant infections.

## INTRODUCTION

The capacity of microorganisms to survive or persist to antibiotics in a nonspecific manner, which is driven by phenotypic diversification rather than by genetic mutagenicity, has been shown to be of relevance in clinical settings [1-5]. Over the years evidence has gradually accumulated which suggests folates are involved in cellular growth regulation. For instance, folinic acid has been observed at high levels (≥70%) in dormant cellular forms, such as plant seeds and fungi spores [6, 7]. Moreover, the overexpression of 5-formyltetrahydrofolate cyclo-ligase (5-FCL), the enzyme that recycles folinic acid has been associated with bacterial dormant phenotypes [8, 9]. Intriguingly, inhibiting 5-FCL in mammalian cell lines affects cell growth [10], and 5-FCL has been described as a pathogenic factor necessary for drug tolerance in *Mycobacterium* [11].

Folate metabolism can be nominally divided into three parts: *de novo* biosynthesis, the folate cycle (equated here to one-carbon folate metabolism or OCFM), and the intake of folate intermediates via facilitated membrane transport [12]. Folate products are involved in anabolic metabolism which is directly coupled with cellular replication activities, such as DNA biosynthesis, and is also associated with the production of methionine, NADPH, and with the transfer of one-carbon groups for epigenetic processes [6, 12-15]. The potential roles of folate metabolism in antibiotic persisters have significant pharmacological applications due to the well established role of antifolates within the pharmacopoeia of antimicrobial and anticancer treatment. Thus, exploring the metabolism of folates could offer venues to intervene and modify the phenotypic traits which facilitate microbial survival to antibiotics.

This study reports measurable levels of antibiotic persisters from the early exponential phase of bacterial growth. The *E. coli* folate gene knockout mutant which is lacking in 5-FCL (*Δfau*) presented lower levels of persisters in comparison to its reference isogenic strain BW25113. Next, we sought to determine if this effect could be observed by exposing fully functional strains to antifolate inhibitors. We found that both BW25113, as well as known hyper-persister (*ΔhipA*) had their survival to bactericidal antibiotics affected. Finally, by adopting an optimised assay for the comparative detection of persisters the gene expression for two different metabolic pathways (including a group of nutrient membrane transporters) was measured. Gene expression profiles had apparent differences among the metabolic states underlying those genetic and pharmacological alterations. These findings contribute to the development of a metabolic framework of the complex cellular states in cell populations where persister phenotypes arise [16].

## METHODS

### Bacterial strains

*E. coli* BW25113 (F-, *Δ(araD-araB)567, Δ lacZ4787* (::rrnB-3), *λ, rph-1, Δ(rhaD-rhaB)568, hsdR514*) is a direct derivative of K12 (BD792) used as the parent strain for the Keio Collection of single gene knockouts [17]. *E. coli* JW2879-1 is isogenic to BW25113 with the gene (*fau* or *ygfA*) encoding 5-FCL deleted (F-, *Δ(araD-araB)567, ΔlacZ4787* (::rrnB-3), *λ, ΔygfA763 ::kan, rph-1, Δ(rhaD-rhaB)568, hsdR514*). *E. coli* JW1500-2 is isogenic to BW25113 with the gene *hipA* deleted (F-, *Δ(araD-araB)567, ΔlacZ4787*(::rrnB-3), *λ, ΔhipA728::kan, rph-1, Δ(rhaD-rhaB)568, hsdR514*). All three strains BW25113, JW2879-1 and JW1500-2 were procured from The Coli Genetic Stock Center. CGSC numbers 7636, 10233 and 9299, respectively.

### Media and Culture

The liquid and solid complex media were Nutrient broth (70122, Sigma-Aldrich, UK) and Nutrient agar (70148, Sigma-Aldrich, UK), respectively. Strains were grown at 37ºC. Where stated M9 media was used as minimal media [18]. Liquid cultures were aerated by shaking in water bath. Every assay reported here started with different fresh cultures from frozen bacterial stocks. Cultures seeded with frozen stocks were incubated typically in 5 mL of culture for 6 to 8 hours under selective antibiotic pressure when appropriate (*Δfau* and *ΔhipA*). These cultures were then diluted 1000-fold in fresh media and incubated overnight. The following day cultures were diluted in fresh media at the same ratio again and used as fresh inocula left to grow up to the desired cell density (OD_600_).

### Antibiotics

Ofloxacin (Sigma O8757), ampicillin (Sigma A9518), and trimethoprim (TMP) (Sigma T7883) were dissolved in H_2_O at stock concentrations of 50 mg/mL. Kanamycin (Sigma K1377) was dissolved in H_2_O at 10 mg/mL, and sulfamethoxazole (SMX) (Sigma S7507) was dissolved in 10 % v/v dimethylsulfoxide (DMSO, Sigma 472301) at 50 mg/mL.

### Growth curves

Fresh inocula were normalised to OD_600_ = 0.01 in fresh broth and cultured in 96-multiwell plates in 0.2 mL. The absorbance at OD_600_ was registered for 24 hours in a Thermo VarioSkan microplate reader (Thermo Fisher Scientific, MA, USA). For the three-hour growth assay overnight cultures were normalised to OD_600_ = 0.1 in 5 mL of fresh broth and aliquoted into as many samples as time points.

### Antibiotic inhibitory concentrations

*E. coli* strains were pre-cultured as detailed in Media and Culture. Final cultures in either minimal or complex media were normalised to OD_600_ = 0.01 by triplicate in 96-well microplates. Antibiotics were present in four-fold dilutions. Growth controls were in triplicate samples without antibiotics. Background controls were included containing 5 mg/L of ofloxacin.

### Three-hour persister assay

Fresh inocula were normalised at OD_600_ = 0.1 in 5 mL of fresh broth. Different samples were withdrawn from incubation at the following time points: 0, 0.25, 0.5, 1, 2 and 3 hours. Then either ampicillin at a final concentration of 100 mg/L or ofloxacin at 5 mg/L were added to each sample and incubated overnight (16-18 hours). All the different samples were then serially diluted 1:10 in 0.2 mL of 10 mM of MgSO_4_ in 96-well microplates. Aliquots of 0.1 to 0.2 mL were plated out on media agar plates and incubated overnight for determination of colony-forming units (cfu) [19].

### Three-hour antifolate assay

Fresh inocula were normalised to OD_600_ = 0.1 in 5 mL of fresh media. Either DMSO (0.1 % v/v), sulfamethoxazole (57.4 mg/L) or trimethoprim (1.7 mg/L) were added, final concentrations within parenthesis. These concentrations correspond to the reported steady-state mean levels of these two antifolates in human plasma during oral administration of Bactrim™ [20]. Samples were withdrawn from incubation at time points (in hours) 0, 0.25, 0.5, 1, 2 and 3. Aliquots of 1.5 mL from each samples were washed twice in fresh complex or minimal media. Samples were normalised to the lowest OD_600_ in 1 mL of media. Then ampicillin at final concentration of 100 mg/L or ofloxacin at 5 mg/L were added to each sample and incubated overnight (16-18 hours). All the different samples were then serially diluted 1:10 in 0.2 mL of fresh media in 96-well microplates. Aliquots of 0.1 mL were plated out on media agar plates and incubated overnight for determination of cfu.

### Real-Time PCR gene expression analysis

Real-Time PCR (RT-PCR) workflow was carried out taking into consideration the MIQE (Minimum information for publication of quantitative real-time PCR experiments) guidelines [21]. The *E. coli* BW25113 genome sequence [22] (GenBank accession number CP009273) was used to design PCR primers (Supplementary Table 1). Three different references genes were used: *cysG, hcaT* and *idnT* [23]. *E. coli* total RNA was extracted with PureZol (7326890, BioRad, UK) following the manufacturer’s recommendations. The quality and quantity of RNA were assessed in the 2100 Bioanalyzer (Agilent Technologies, Santa Cruz, CA). Synthesis of cDNA was performed with iScript (Biorad, UK) in 0.02 mL reactions scaled up when required up to 0.12 mL. Amplification reactions were set up in 0.01 mL using SsoAdvanced Universal SYBR Green Supermix (Biorad, UK) following manufacturer’s recommendations. PCR reactions were distributed in Hard Shell low profile skirted 96-well microplates sealed with Microseal B films (Biorad, UK). Amplification reactions (0.01 mL) were carried out in a CFX96 Real-Time System thermocycler (Biorad, UK), with the results compiled and automatically analysed at source with the CFX Manager software (Biorad, UK). The data presented represent three different biological replicates (n=3). Plated samples also included eight 6-fold dilutions of cDNA, with a known amount of DNA and using primers for *cysG*, for the generation of a standard curve. This allowed quantitation of gene amplification within each plate. Amplification efficiency was measured to assess the reproducibility of the amplification for each gene. Quantitative comparative gene amplifications was carried with normalised detection values and used to generate the fold changes as log ratios with change p-values (Supplementary Tables 14 - 16). The analysed data were compiled and plotted as heatmaps using heatmap2, which is a part of R’s gplots [24].

**Table 1.** The inhibitory concentrations of antibiotics and the solvent (DMSO) used in this study. All three strains of *E. coli* are included. BW25113 is the parental strain, *Δfau* is the folate mutant strain, and *ΔhipA* is the hyper-persister strain. Data represent inhibitory concentrations as IC_50_ in mg/L, except for DMSO which is in % (vol/vol).

|  | Minimal media |  |  | Complex media |  |  |  |
| --- | --- | --- | --- | --- | --- | --- | --- |
|  | BW25113 | *Δfau* | *ΔhipA* | BW25113 | *Δfau* | *ΔhipA* | Used |
| DMSO | 5.7 ± 0.67 | 13.7 ± 0.47 | 6.62 ± 0.41 | 8.173 ± 1.743 | 5.746 ± 0.952 | 7.166 ± 1.98 | 0.1 |
| Ampicillin | 3.96 ± 0.3 | 3.93 ± 0.423 | 2.31 ± 0.017 | 6.61 ± 0.718 | 5.644 ± 0.563 | 4.969 ± 0.499 | 100 |
| Ofloxacin | 0.234 ± 0.004 | 0.234 ± 0.0043 | 0.249 ± 0.003 | 0.084 ± 0.023 | 0.095 ± 0.015 | 0.088 ± 0.007 | 5 |
| Trimethoprim | 0.59 ± 0.027 | 0.539 ± 0.022 | 0.62 ± 0.011 | 0.572 ± 0.082 | 1.064 ± 0.242 | 0.899 ± 0.138 | 1.7 |
| Sulfamethoxazole | 0.39 ± 0.023 | 0.132 ± 0.01 | 0.464 ± 0.032 | 349 ± 8.55 | 141 ± 2.83 | 2383 ± 262 | 57.4 |

### Statistical analysis

Numerical analysis and graphical production were performed using R [24]. Antibiotic inhibitory concentrations were calculated with the dose-response analysis R extension package *drc* [25]. Principal Components Analysis (PCA) was carried out using the R packages FactoMineR and Factoextra [26]

## RESULTS

Persistence to antibiotics is a phenotypic stochastic phenomenon that arises at low frequency in fresh media [27]. However, certain genetic backgrounds, such as the hyper-persister toxin *hipA* mutant [28, 29] or the altered gene expression of certain genes [8] have been shown to influence the frequency of persisters. An example of the latter is the decrease in the rate of persisters under low expression of the gene that encodes for the folate enzyme 5-FCL [8]. The *E. coli* gene knockout strains *Δfau* and *ΔhipA* and their isogenic parental strain (BW25113) were used throughout this work. Thus, we first measured the sensitivity of these three strains to each of the different antibiotics used in this study in both complex and minimal media. Two significant differences were found, the expected increased sensitivity of the folate knockout strain *Δfau* to SMX in the minimal media, and the high inhibitory concentrations (IC_50_) of SMX in complex media (Table 1, Supplementary Figs. 1 and 2).

Importantly, the concentrations of ampicillin and ofloxacin used for the selection of persisters were above the IC_50_ observed for these strains. Also, the concentrations of antifolates (with the exception of SMX in complex media) used here were above the inhibitory concentrations (Table 1). Significantly, the inhibitory concentrations of antifolates in vitro were not decisive for the concentrations of antifolate to use in the persister assays. Our prerogative was to assess concentrations in line with to the steady-state levels found in human plasma during treatment with oral formulation of SMX and TMP [20].

### Growth rates

In contrast to previous investigations [8, 28, 29] it was found that the growth rate in *Δfau* and *ΔhipA* was different to the parental strain, BW25113, in a time and media-dependent fashion. Cultures followed for 24 hours showed differential growth rates between the three strains used here, particularly in the complex media (Fig. 1). Therefore, it was deemed necessary to establish incubation times that allowed observations to be made without visible differences in growth rate. We also were primarily interested in studying persisters arisen under non apparent stress (i.e., cells in logarithmic growth in fresh media), also known as Type II persisters [3]. Phenotypic diversification present at early moments of cell growth can be relevant to persister formation in the logarithmic growth phase [30]. Based on these premises, batch cultures were followed throughout 3 hours with a starting OD_600_ = 0.1, optimised to allow the retrieval of cells growing logarithmically in either complex or minimal media (Fig. 2). Noticeably, the generation time approximately doubled in the minimal media (1.5 ± 0.13 hours), in comparison to bacteria which grew in the complex media (0.66 ± 0.07 hours) for all three strains (Figs. 1 and 2). Importantly, within this time window and at the observed cell densities no differences in growth rates among strains were observed independent of the type of media and the cell counts per unit of OD_600_ were comparable among strains, BW25113 (95% CI 4.9 - 5.1 x 10^8^ cells), *Δfau* (95% CI 4.9 - 5.2 x 10^8^ cells) and *ΔhipA* (95% CI 5.0 - 5.15 x 10^8^cells).

**Fig. 1.**
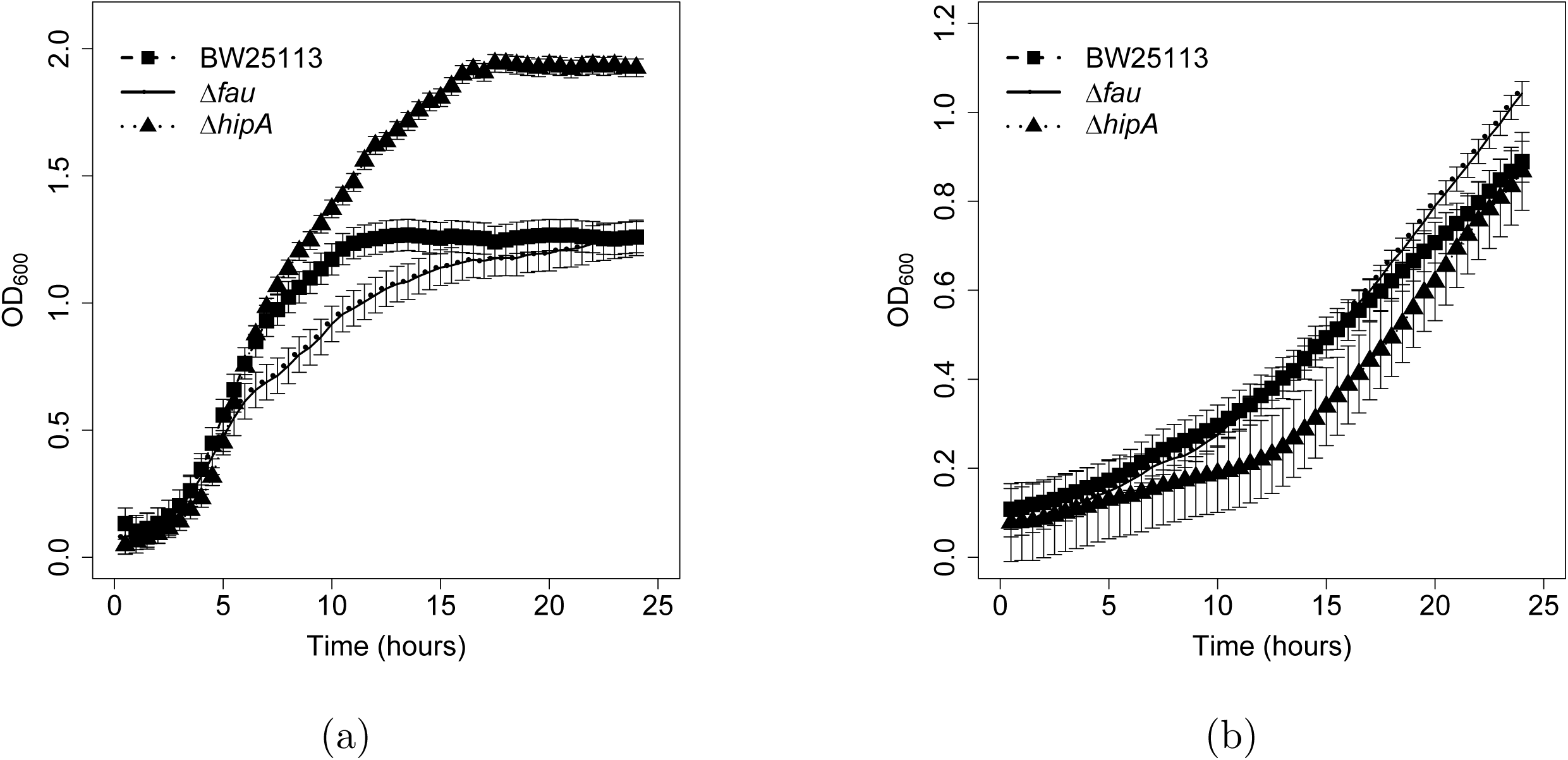
Growth rates (24 hours). Bacterial growth was followed for 24 hours in either complex media (a) or minimal media (b). The Y-axis represents the optical density (OD_600_) with a starting OD_600_ = 0.01. The means and standard deviations for three different experiments (n = 3) are shown. BW25113 denotes the reference strain, and *Δfau* and *ΔhipA* the gene knockout strains for the folate gene *fau* and the toxin gene *hipA*, respectively.

**Fig. 2.**
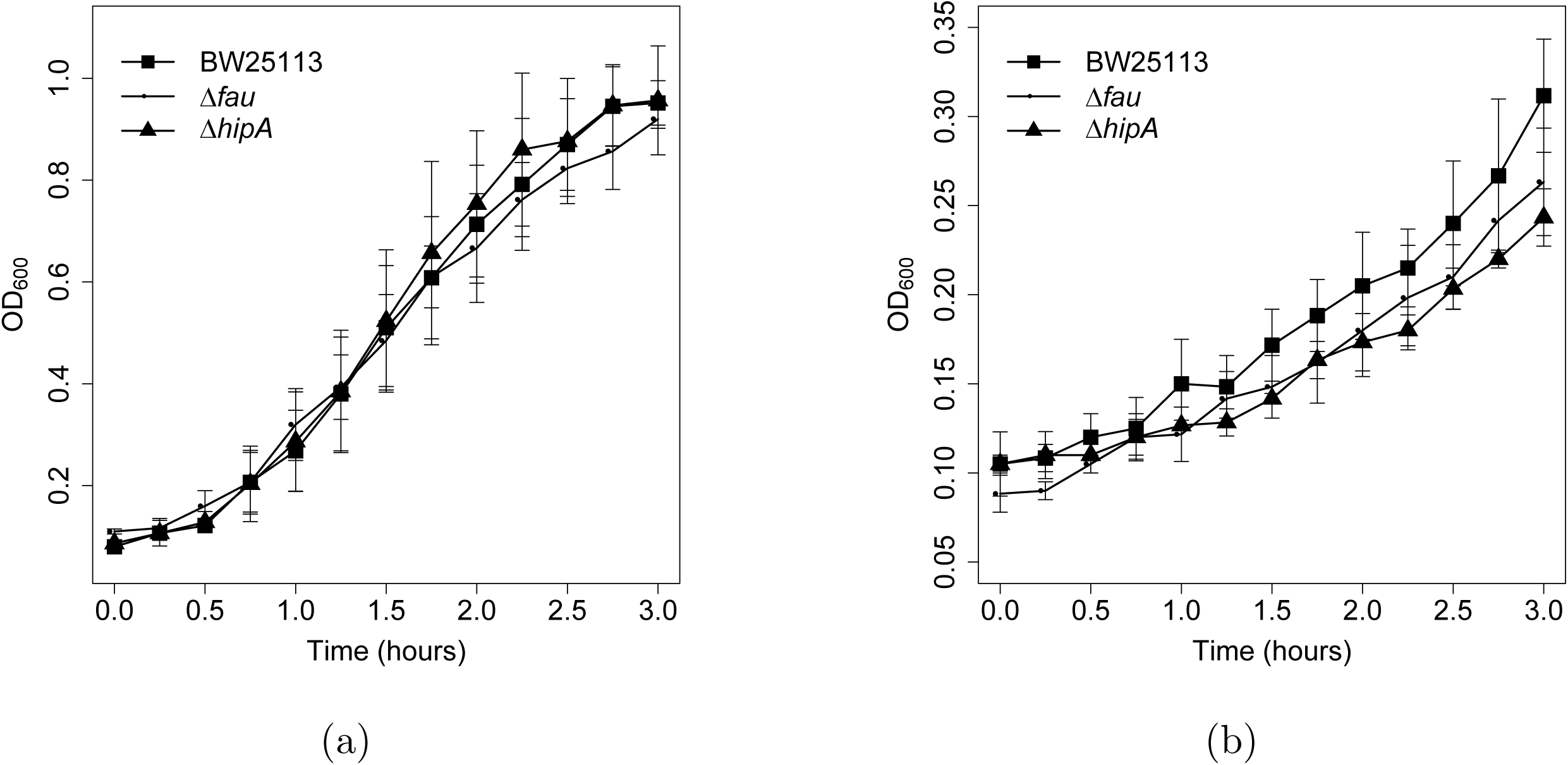
Growth rate (3 hours). Bacterial growth was followed for 3 hours in either complex media (a) or minimal media (b). The Y-axis represents the optical density (OD_600_) with a starting OD_600_ = 0.1. The mean and standard deviation for three different experiments (n = 3) are shown.

### Biphasic response to ampicillin and ofloxacin

Low-level-persistence phenotypes are usually observed as biphasic responses to antibiotics. At a OD_600_ = 0.3, both the reference and the folate mutant strains displayed a time-dependent killing level of persisters which stabilised after 6 hours of incubation in ampicillin in either complex or minimal media (Fig. 3). The folate mutant continued to show a decrease of persisters to ampicillin in complex media after 6 hours. At 16 hours we measured the incidence of persisters to ampicillin at 95% CI 320 - 400 cfu per 10^8^ cells for BW25113 and at 95% CI 56 - 80 per 10^8^ cells for *Δfau*. A similar pattern but fewer persisters was seen for ofloxacin in BW25113 (95% CI 8 - 20 per 10^8^ cells) and *Δfau* (95% CI 4 - 8 per 10^8^ cells).

**Fig. 3.**
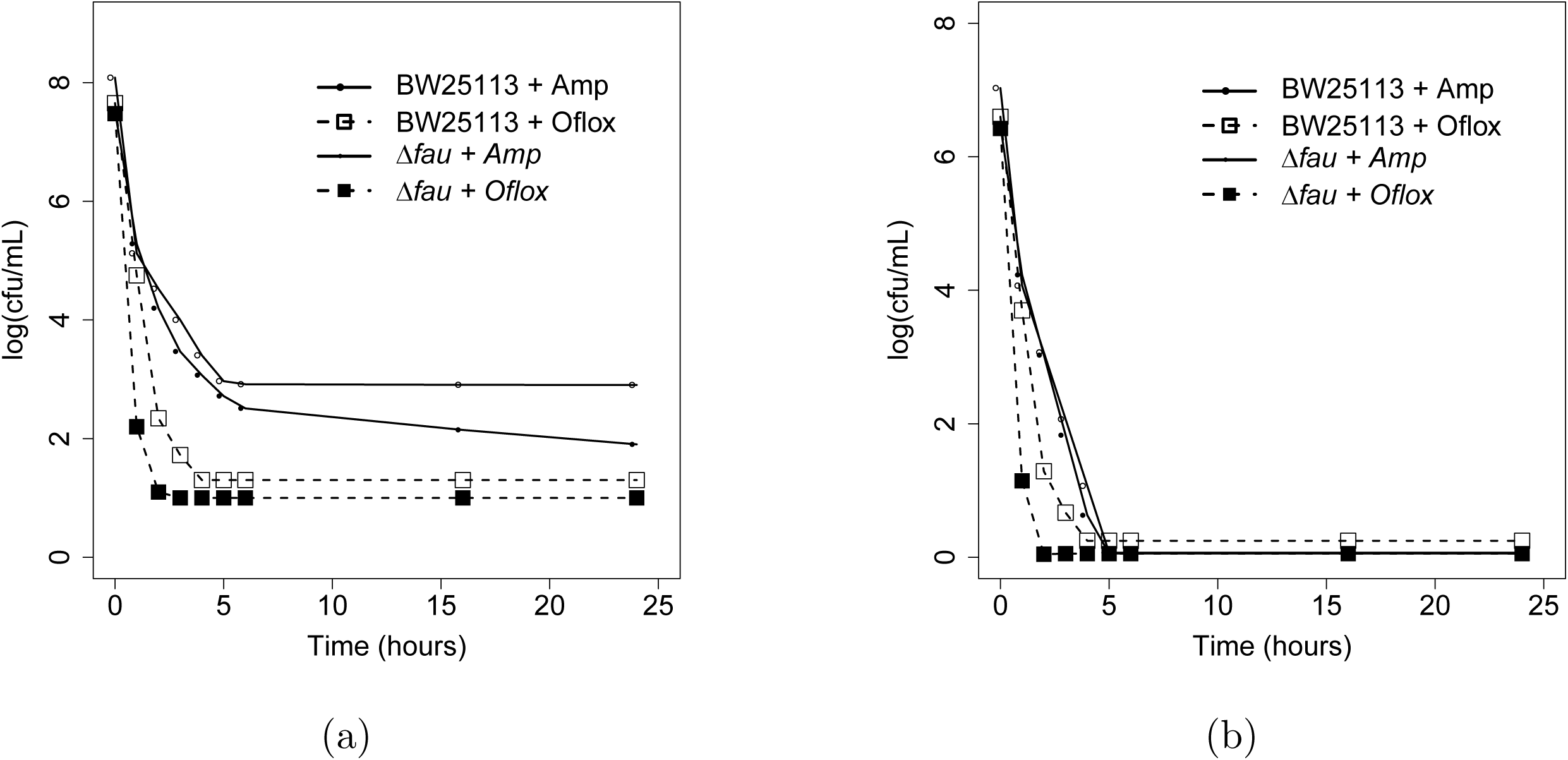
Biphasic time-dependent killing. Cultures were grown to an OD_600_ = 0.3 before adding either ampicillin (100 mg/L) or ofloxacin (5 mg/L) in complex media (a) or minimal medium (b) for the indicated times. Samples collected at a given time were diluted and spread plated on LB agar. The experiments were performed in triplicate (n=3).

### Rate of persistence in cells in logarithmic growth (the 3-hour assay)

The appearance rate of persisters is presented as ratios of cfu/mL from a given batch culture at each time over the cfu/mL of the batch culture at time zero. This normalises the persister phenotypes related to the physiological status of the inoculates at the starting point. By three hours in complex media BW25113 had twice more persisters to ampicillin than *Δfau* (Fig. 4a). BW25113 presented again more persisters to ofloxacin than *Δfau*, this time by near a five-fold difference (Fig. 4b). Remarkably, the number of persisters for either strain increased consistently from their lag phase and along their growth in exponential phase.

**Fig. 4.**
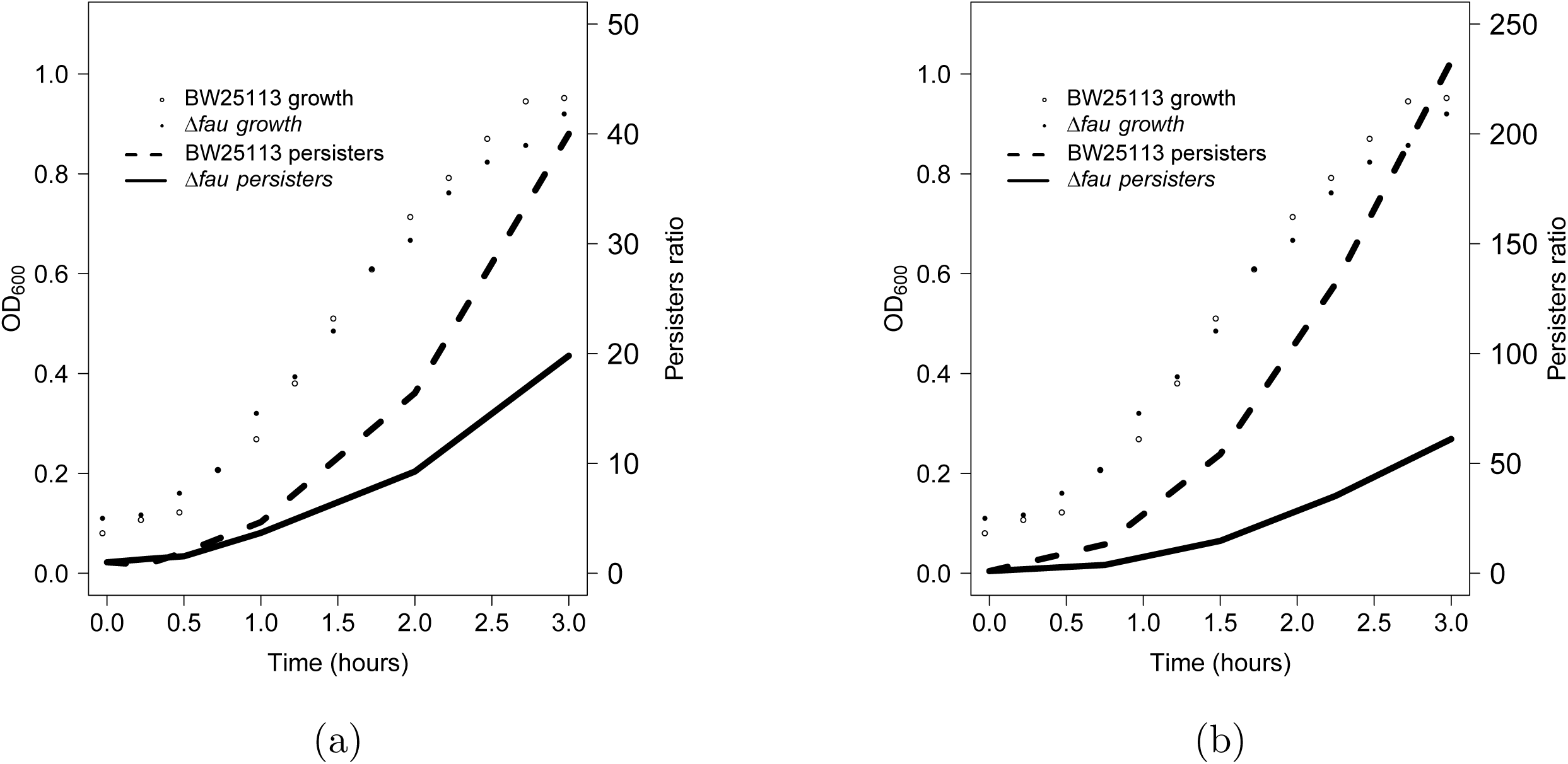
The incidence of persisters in the folate gene knockout strain in complex media. Bacteria were exposed for 16 hours to ampicillin 100 mg/L (a) or ofloxacin 5 mg/L (b), after growing in complex media for different durations (0, 0.25, 0.5, 1, 2 and 3 hours). The left Y-axis shows the optical density of the cultures and the right Y-axis shows the ratios (versus time zero) of persisters after two hours of incubation in either antibiotic. Data compiled in the Supplementary Tables 1 and 2.

In sharp contrast, the ratio of persisters in the minimal media for BW25113 and *Δfau* declined throughout, and were closed to zero by 3 hours (Fig. 5). This applied to both, bacteria exposed to ampicillin and ofloxacin. The metabolic network of cells in minimal media was clearly incompatible with the necessary metabolic state required for the phenotypic variations driving persistence.

**Fig. 5.**
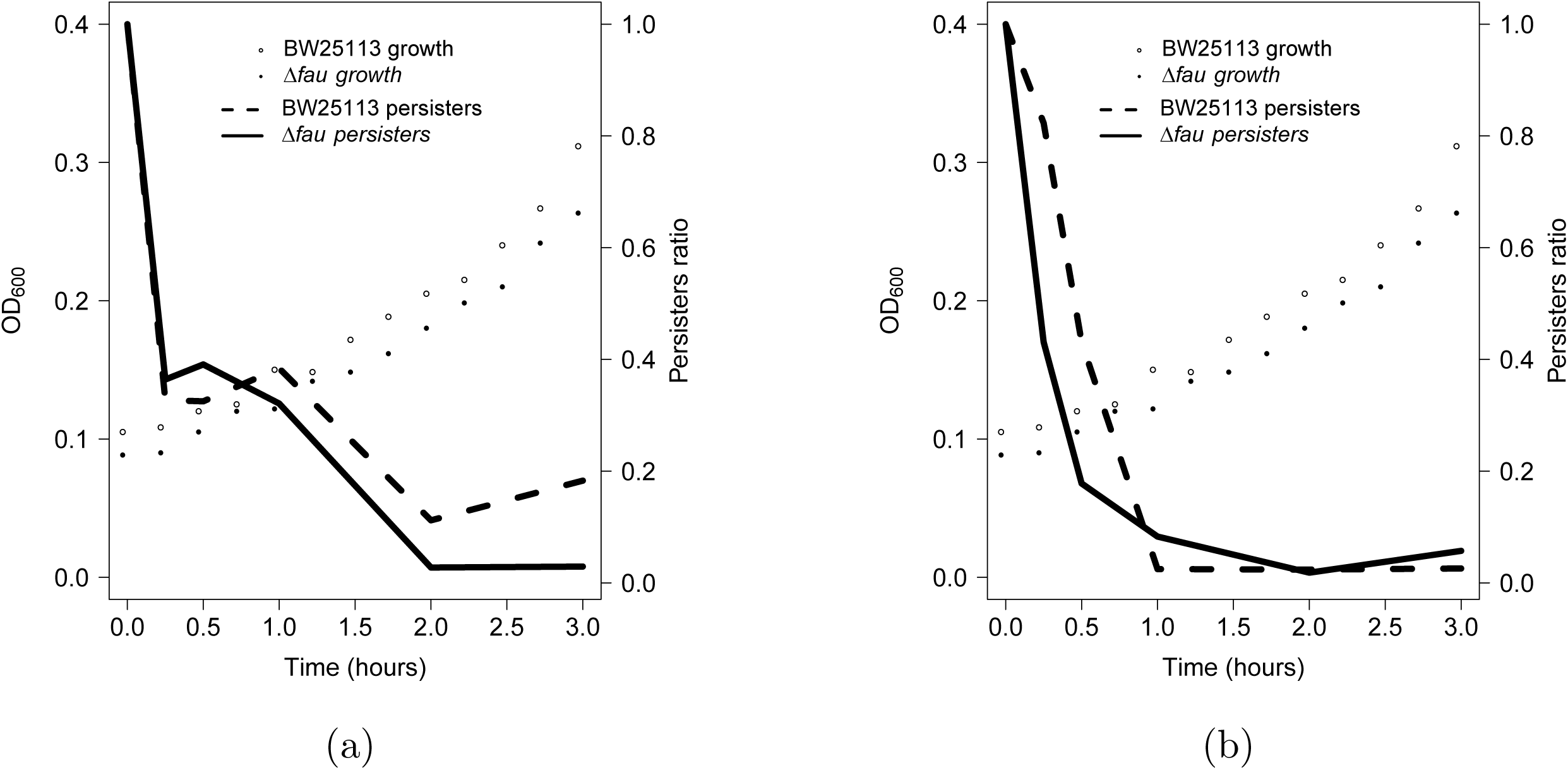
The incidence of persisters in the folate gene knockout strain in minimal media. Bacteria in minimal media were exposed, at different time points (0, 0.25, 0.5, 1, 2 and 3 hours), to ampicillin 100 mg/L (a) or ofloxacin 5 mg/L (b). The left Y-axis shows the optical density of the cultures and the right Y-axis shows the the ratios (versus time zero) of persisters after 16 hours of incubation in either antibiotic. Data compiled in the Supplementary Tables 3 and 4.

### The effect of antifolates on the reference strain BW25113

It was considered relevant to study how the results presented thus far compared with the behaviour of the bacteria following administration of pharmacological inhibitors of folate biosynthesis, and how this affected the development of antibiotic persisters. In the absence of validated inhibitors of 5-FCL it was deemed cogent to use two well established antimicrobial antifolates: sulfamethoxazole (SMX) and trimethoprim (TMP). Inhibitors of the de novo folate biosynthesis enzyme dihydropteorate synthetase, and the OCFM enzyme dihydrofolate reductase, respectively. The BW25113 strain was incubated for up to three hours using each of these antifolates or DMSO as the control for the solvent used for SMX. As expected, growth was affected by TMP in the complex media and by both SMX and TMP in the minimal media (Figs. 6a and 6d). Therefore, samples had to be normalised based on their optical densities to the sample with the least cell mass per volume. Thereafter, cultures were exposed overnight to either ampicillin or ofloxacin.

**Fig. 6.**
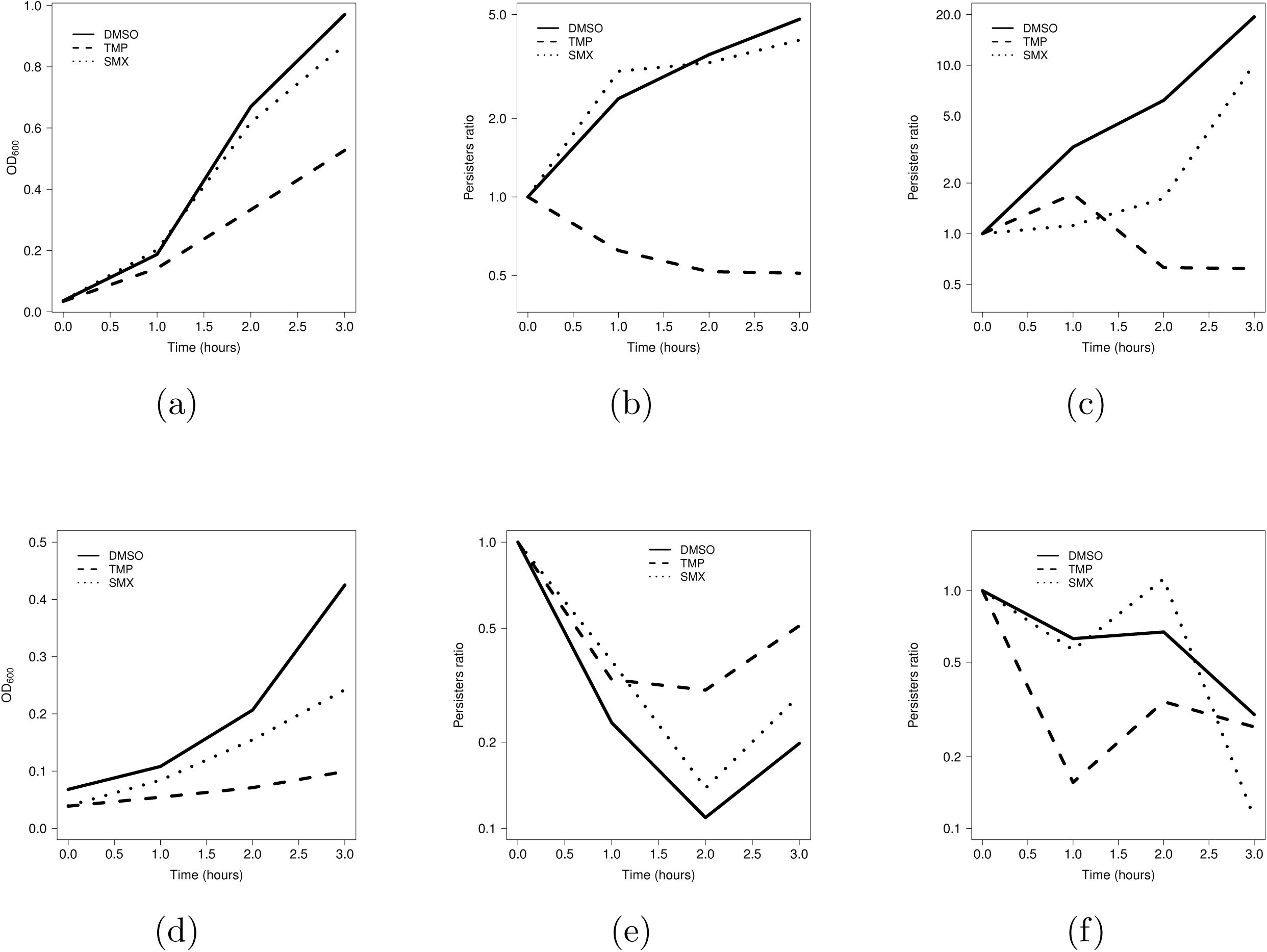
The effects of antifolates on persistence to antibiotics in the reference strain BW25113. (a) Bacteria in complex media in the presence of either 0.1% (v/v) dimethylsulfoxide (DMSO), 1.7 mg/L trimethoprim (TMP), or 57.4 mg/L sulfamethoxazole (SMX). At different time points (0, 1, 2, and 3 hours) the cultures were washed, resuspended in complex media and incubated for 16 hours in the presence of either antibiotic, ampicillin 100 mg/L (b) or ofloxacin 5 mg/L (c). Incidence of persister is shown as a ratio of cfu/mL from each time point over time zero. (d) Bacteria in minimal media in the presence of either 0.1% (v/v) dimethylsulfoxide (DMSO), 1.7 mg/L trimethoprim (TMP), or 57.4 mg/L sulfamethoxazole (SMX). At different time points (0, 1, 2, and 3 hours) the cultures were washed, resuspended in minimal media and incubated for 16 hours in the presence of ampicillin 100 mg/L (e) or ofloxacin 5 mg/L (f). Incidence of persister is shown as a ratio of cfu/mL from each time point over time zero. Data for Figs. 6b and 6c compiled in Supplementary Tables 5 and 6. Data for Figs. 6e and 6f compiled in Supplementary Tables 7 and 8.

Incubation with TMP in complex media clearly repressed the development of persisters to ampicillin throughout and from the two-hour point for ofloxacin (Figs. 6b and 6c). The effect of SMX is not apparent for ampicillin but interestingly it delayed the appearance of persisters to ofloxacin after two hours (Fig. 6c). The metabolism of bacteria in minimal media was still affected by the incubation in TMP given the higher ratio of persister for ampicillin than in the solvent control and SMX (Fig. 6e). However, the ratio of persisters for ofloxacin fluctuated in all three samples, from incubations with the two antifolates and DMSO, and by the third hour none had more persisters than at time zero (Fig. 6f).

### The effect of antifolates on the hyper-persistent mutant strain *ΔhipA*

By the very nature of random phenotypic diversification microbes are likely to be in a broad spectrum of physiological states while growing logarithmically. It was reasoned important to investigate the response to antifolates from a strain prone to develop persisters by using the knockout mutant of toxin hipA (*ΔhipA*), a well known hyper-persister [28, 29]. In contrast to SMX, TMP had a discernible inhibition on the growth of *ΔhipA* (Fig. 7a). Subsequently, TMP showed to have a clear suppressing effect on the ratio of persisters in *ΔhipA* in complex media for both ampicillin and ofloxacin (Figs. 7b and 7c). In a trend not dissimilar to the effects on the parental strain (Figs. 6b and 6c). In minimal media both SMX and TMP reduced the growth rate of *ΔhipA* (Fig. 7d). Both antifolates did not alter the patterns observed for the parental strain as consistently lower ratios of persisters in this type of media were observed for ampicillin in all samples after time point zero (Fig. 7e). For ofloxacin both antifolates had an effect in *ΔhipA* more apparent than in the parental strain and were significantly different to the effect of DMSO (Fig. 7f).

**Fig. 7.**
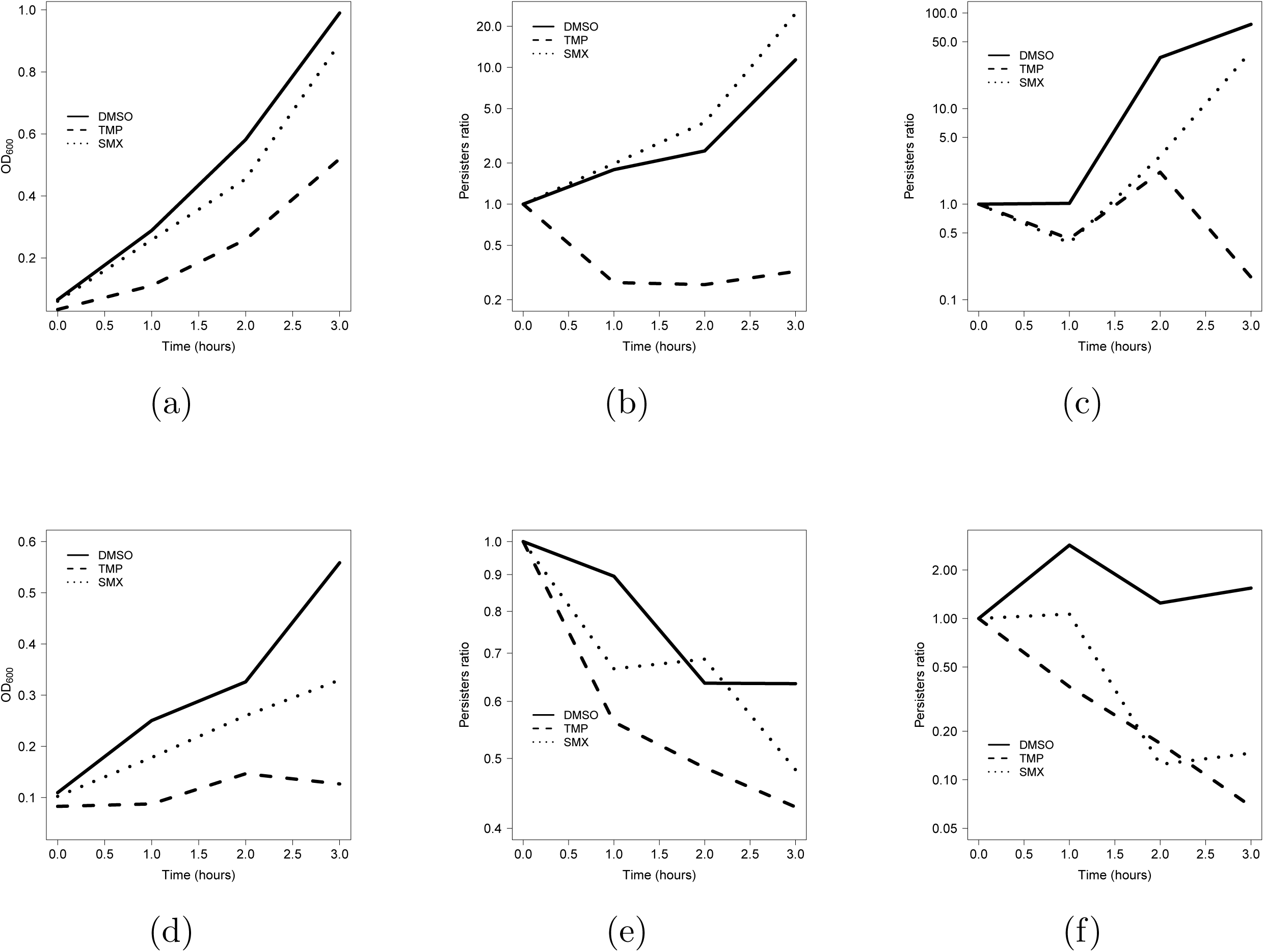
The effects of antifolates on the persistence to antibiotics in the hyper-persister strain *ΔhipA*. (a) Bacteria in complex media in the presence of either 0.1% (v/v) dimethylsulfoxide (DMSO), 1.7 mg/L trimethoprim (TMP), or 57.4 mg/L sulfamethoxazole (SMX). At different time points (0, 1, 2, and 3 hours) the cultures were washed, resuspended in complex media and incubated for 16 hours in the presence of either antibiotic, ampicillin 100 mg/L (b) or ofloxacin 5 mg/L (c). The incidence of persisters is shown as a ratio of cfu/mL from each time point over time zero. (d) Bacteria in minimal media in the presence of either 0.1% (v/v) dimethylsulfoxide (DMSO), 1.7 mg/L trimethoprim (TMP), or 57.4 mg/L sulfamethoxazole (SMX). At different time points (0, 1, 2, and 3 hours) the cultures were washed, resuspended in minimal media and incubated for 16 hours in the presence of ampicillin 100 mg/L (e) or ofloxacin 5 mg/L (f). The incidence of persisters is shown as a ratio of cfu/mL from each time point over time zero. Data for Figs. 7b and 7c compiled in the Supplementary Tables 9 and 10. Data for Figs. 7e and 7f compiled in the Supplementary Tables 11 and 12.

The control (DMSO) was included in the pharmacological assays with antifolates at a final concentration equivalent to that in SMX (0.1% v/v). Unexpectedly, samples with DMSO at this final concentration did not necessarily follow the behaviour of the samples with SMX. This solvent inhibited the development of persisters for ampicillin in the minimal media in both the parental strain BW25113 and the hyper-perister *ΔhipA* (Figs. 6e and 7e). A trend not observed for ofloxacin in the same media (Figs. 6f and 7f), nor for ampicillin or ofloxacin in the complex media (Figs. 6b, 6c, 7b, and 7c).

### The impact of Antifolates impact on persister development: Principal Components Analysis

Principal Components Analysis (PCA) was used to summarise the data gathered for the effects of antifolates BW25113 and *ΔhipA*. Data were organised as listed in the Supplementary Table 13: presence of antifolates, type of antibiotic (ampicillin or ofloxacin), strain (BW25113 or *ΔhipA*), time spent in antifolates (0 - 3 hours), media (complex or minimal), and ratio of persisters (cfu/mL at a given time up to 3 hours over cfu/mL at time zero). The first two dimensions made up approximately 41% of the total variance of the dataset (inertia) (Fig. 8). The variables type of antibiotic and time spent in antifolates were correlated positively to the ratio of persisters. This was a key finding as it suggested persister development was associated with the type of antibiotic (ampicillin or ofloxacin) in agreement with the literature, but also here in relation to the time spent in the antifolates. The variables media and the type of antifolates were negatively correlated to the ratio of persisters, indicating that in either media either antifolate seemed to lower the incidence of persisters (Fig. 8). Notably, the variable strain, the reference strain BW25113 or the hyper-persister *ΔhipA*, behaved as a supplementary (*i.e.,* non-active) variable. This suggested antifolates had a similar effect on both of these strains. That is to say the hyper-persister strain was similarly susceptible to the effects of antifolates. An encouraging finding for clinically relevant settings.

**Fig. 8.**
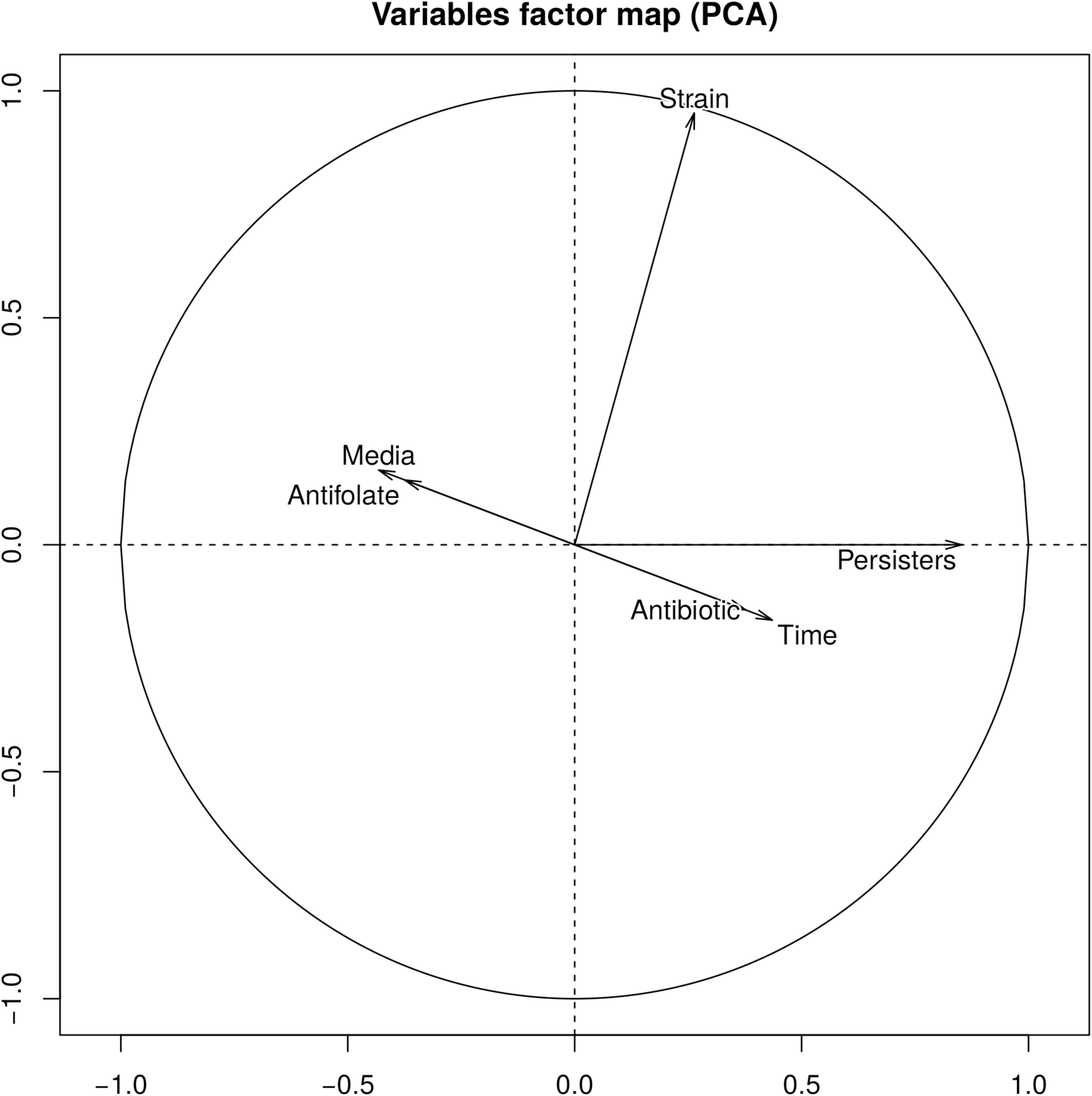
Principal Components Analysis for the effect of antifolates on the development of persisters. The dataset on antifolates was summarised with six different variables: strain (BW25113 or *ΔhipA*), time exposure to antifolates (in hours), antifolate (DMSO, SMX, TMP), media (minimal media or complex media), antibiotic (ampicillin or ofloxacin), and level of persisters (the ratio of cfu/mL at each time point in antifolates over the cfu/mL at time zero). The X-axis represents the principal component 1, and the Y-axis represents the principal component 2. Distances from zero represent the magnitudes of the effects of each variable. The associated ranks in order to carried out the PCA are in Supplementary Table 13.

### The gene expression profile of the reference strain BW25113 in exponential growth

It was logical to investigate the molecular basis for the development of persisters in the reference *E. coli* strain. A methodology was developed to use optimised targeted RT-PCR amplifications to profile the gene expression responses of the parental *E. coli* strain, BW25113 throughout the 3-hour growth assay in the complex media (Fig. 2a). Cells that grew within this time frame displayed the persistence phenotypes associated with both ampicillin and ofloxacin (Fig. 4). The set of genes which were queried fully encode for folate biosynthesis and usage (OCFM) pathways, fifteen solute transporters, twelve fatty acids biosynthesis enzymes, and the *cydX* gene (subunit X of cytochrome d (bd-I) ubiquinol oxidase) (Supplementary Table 14). Also included were fatty acid biosynthesis genes which have been shown to be involved in the development of persisters [31]. On the other hand, oxidative phosphorylation, represented here by *cydX*, has been shown to sensitise bacteria to antibiotics [32, 33]. Consistently, cydX was significantly underexpressed in growth phases where bacteria produced persisters to antibiotics (Fig. 9). Data were analysed and interpreted by comparing the gene expression levels at different time points against the initial samples (time zero). Folate biosynthesis represented by *folC, pabC, aroH, aroC*, and *folP* appeared to be active through the initial hour of growth in fresh complex media (Fig. 8). Only the transport of amino acids and pantothenate, *putP* and *panF*, were overrepresented before the initial hour. After the first hour, which represents a cell mass equivalent to an OD_600_ = 0.3 (Figs. 2a and 4), these metabolic components were clustered with genes representing other enzymes of folate biosynthesis as well as nine more substrate membrane transporters (*cycA, shiA, citT, nupC, dsdX, nupG, aroP, glcA*, and *uhpT*) (Fig. 9).

**Fig. 9.**
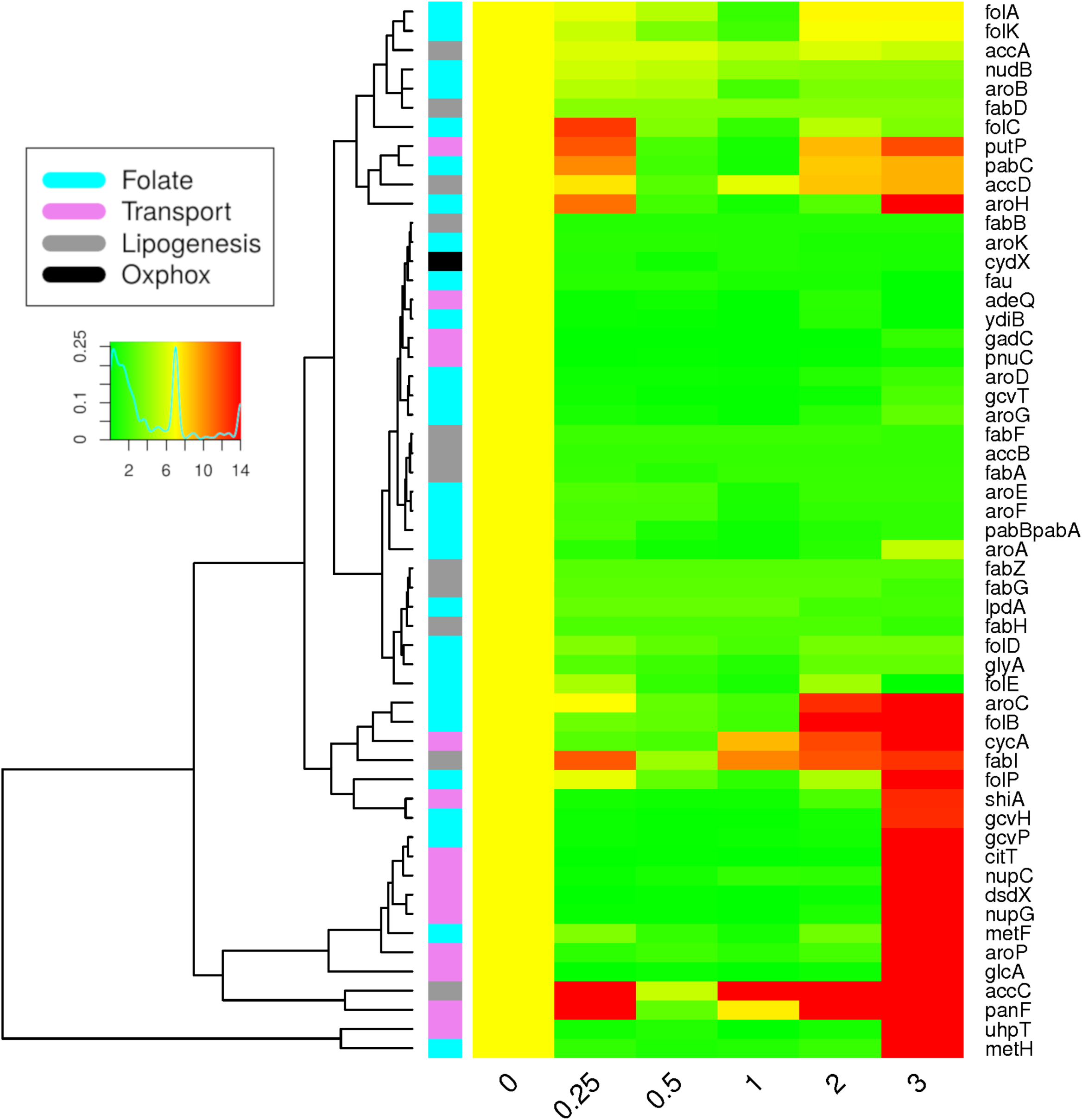
Gene expression in the reference strain (BW25113) during the three-hour growth assay in complex media. The levels of gene expression, at the specified time points (in hours), as a ratio of the values at time zero. The yellow column at the initial time point denotes no differences in gene expression since this the ratio of gene expression levels in this sample against itself. Colour keys for the metabolic pathways which were studied are to the left of the heatmap (respiration gene *cydX* denoted by the black colour key) and the gene list is to the right.

### The comparative gene expression profile of the folate mutant strain *Δfau* in exponential growth

We assessed the genetic expression profile of the gene knockout folate mutant *Δfau* in comparison to BW25113. The initial outline at time zero showed the majority of genes underexpressed with three genes overexpressed in *Δfau* cells, all three solute membrane transporters (*gadC, uhpT, citT*) involved in the uptake of organic acids and carbohydrates (Fig. 10). During the first hour there was an apparent overexpression of genes that included the glycine cleavage complex (GCV) of the OCFM (*lpdA, gcvH, gcvP, gcvT*), whose gene expression pattern clustered with the proline:Na symporter *putP*. The time points at two and three hours (Fig. 10) gave an indication of the potential the overexpression of a number of genes involved in encoding lipogenesis enzymes. The majority of solute membrane transporters targeted here were overexpressed by the second or third hour (*shiA, nupG, gadC, nupC, cycA, panF, uhpT, glcA, aroP, citT, adeQ*, and *dsdX*). Altogether, folate metabolism in the folate mutants was underrepresented, while lipogenesis, and a number of solute membrane transporters seem to have been overrepresented.

**Fig. 10.**
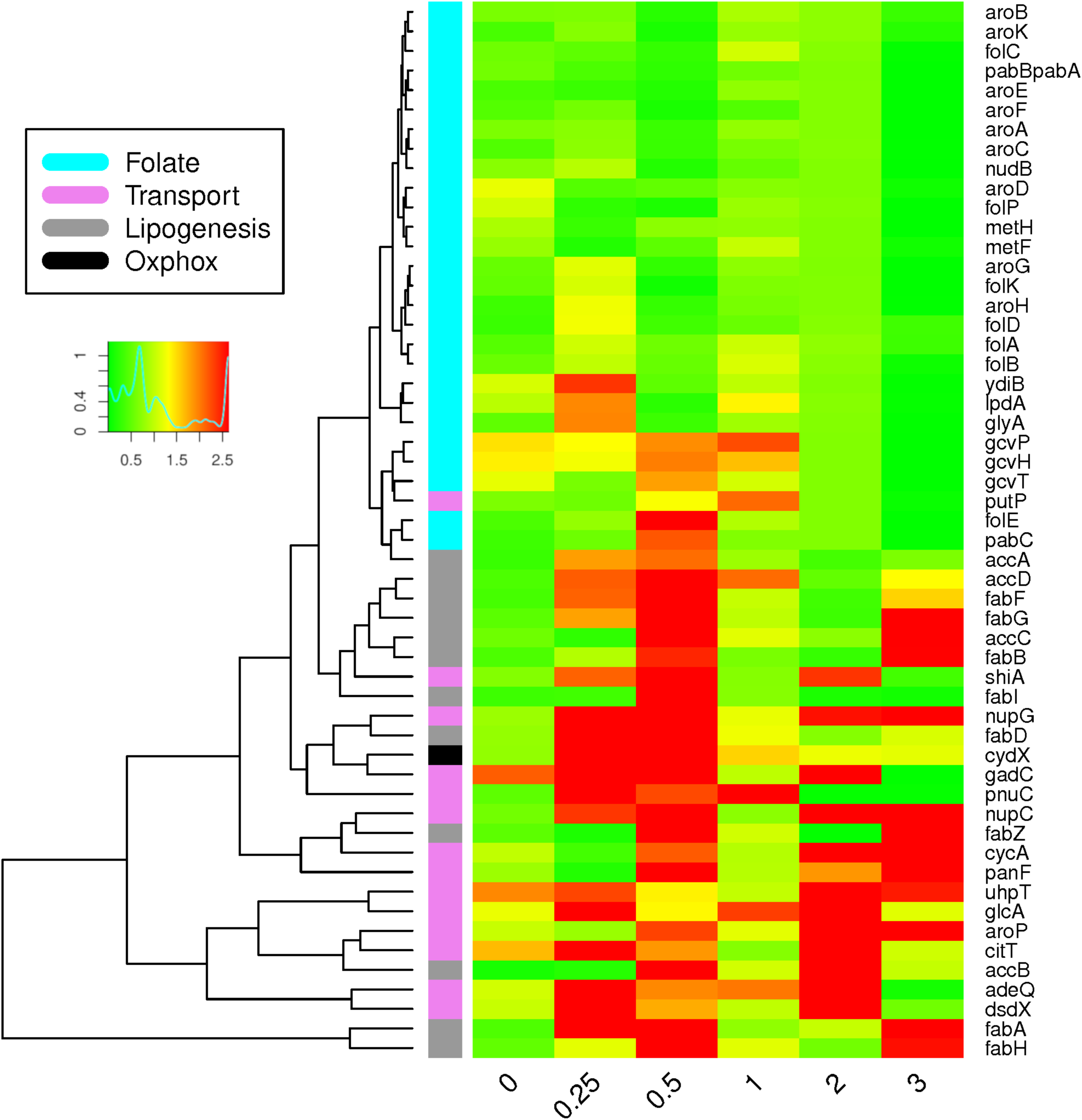
Comparative differential gene expression in the folate mutant strain. The levels of gene expression, at the specified time points (in hours), in the folate knockout mutant *Δfau* strain compared to the reference strain BW25113. Colour keys for the metabolic pathways which were studied are to the left of the heatmap (respiration gene *cydX* denoted by the black colour key) and the gene list is to the right.

### The comparative gene expression profile of the bacteria exposed to trimethoprim

In the complex media only TMP was found to alter the incidence of persisters to ampicillin and more significantly to ofloxacin (Fig. 5). BW25113 cells treated with TMP had initially a discrete number of overexpressed genes encoding folate metabolism and lipogenesis together with two membrane transporters (*aroP* and *adeQ*) (Fig. 11). After one hour, the over-expression of genes encoding for folate metabolism (particularly representing OCFM) and lipogenesis became more apparent (Fig. 11). Similarly, five solute membrane transporters were over-expressed after the first hour (*cycA, pnuC, panF, adeQ*, and *nupG*). Contrary to the *Δfau* gene knockout strain, the response to antifolate treatment seems to have included a representation of folate biosynthesis (Figs. 9 - 11).

**Fig. 11.**
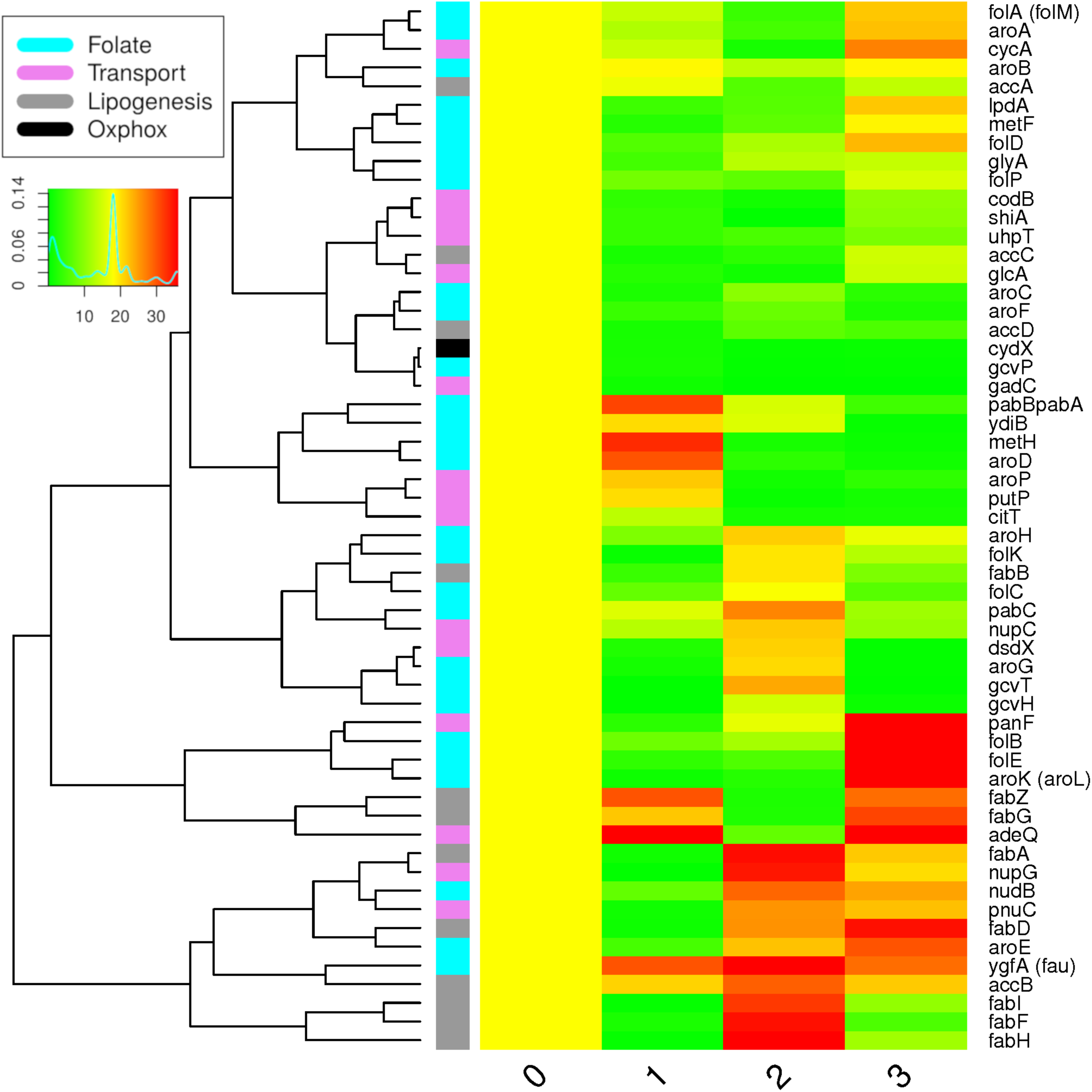
Gene expression in the reference strain (BW25113) in the presence of trimethoprim. Gene expression, at the specified time points (in hours), in the BW25113 strain incubated in trimethoprim (1.7 mg/L) when compared to the untreated BW25113 control. The yellow column at the initial time point denotes no differences in gene expression at the moment of adding the antifolate. Colour keys for the metabolic pathways which were studied are to the left of the heatmap (respiration gene *cydX* denoted by the black colour key) and the gene list is to the right.

## DISCUSSION

The evidence presented here connects folate metabolism to the phenotypes underlying the survival of exponentially growing bacteria exposed to antibiotics. Although phenotypic variation is expectedly based on the known cell to cell diversification and single cell gene to gene expression variability [34], it is unknown how this diversity allows sub-populations of microorganisms to survive environmental stressors and qualified as persisters. In an attempt to conceptualise these phenomena some have called persistence a consequence of dormancy or latency. Here, we are avoiding these terms which may not necessarily convey the physiology of cells in rapid growth that happen to survive a given stressor [16, 35].

Antibiotic peristers were described soon after the discovery of penicillin [36] and subsequently observed across different forms of drug tolerance in human pathogens [3, 4, 37]. Nonetheless, bacteria surviving exposure to a stressor could be simply a by-product of the events preceding their formation. The biochemical composition of the bacterial cell prior to exposure which enables a numbers of the population to remain viable and to successfully reproduce is the conundrum which needs to be solved. In some scenarios, such as cells in apparent nutritional stress (i.e., bacteria in spent media) the development of persisters to antibiotics is three to four orders of magnitude higher than in a population in fresh media [38, 39]. This has been rationalised under the known cell signalling that takes place under nutritional stress [40]. However, the study of how persisters appear in cell growing exponentially may not benefit from the same rationale. Bacteria viable after exposure to antibiotics are observed before they reach stationary growth (Type II persisters) [41-42].

We observed that persisters to antibiotics are detectable from the very moment when bacteria leave the lag phase in complex media (Fig. 4). Strikingly, the development of persisters is rapid along logarithmic growth (Fig. 4). Therefore, part of the information needed to reconcile these events could be in the very nature of exponential growth. For instance, there is measurable cell to cell heterogeneity (e.g., cell size variability, DNA content, protein content) intrinsic to rapid cell growth [43-44].

Rapid cell growth and the generation of phenotypes that facilitate the formation of persisters could be two sides of the same coin [42]. Different dynamics (i.e., pathway specific fluxes) of anabolic metabolism would be expected to be part of the phenotypes which allow persistence to antibiotics. Consistent with this, batch cultures in the 3-hour assays were observed in the complex media to have a significant proportion of persisters to ampicillin and ofloxacin (Fig. 4), while bacteria in the minimal media at equivalent cell mass showed no detectable persisters (Fig. 5). The gene expression profile in complex media for the reference strain showed clusters of overexpression that by the third hour had possibly 25 overexpressed genes, for folate and lipid biosynthesis, in comparison to the beginning of the lag phase (Fig. 9).

The anabolic roles of folate as a major source of carbon for DNA biosynthesis, methionine biosynthesis, and NADPH production, also include folate as a critical carbon donor in epigenetic modifications, including those observed in prokaryotes [14]. It can be speculated that a role of folate which impacts the persister phenotype could include epigenetic changes, together with its more established role in anabolic metabolism. The idea of persister phenotypes driven by epigenetic changes has been proposed by *in silico* modelling [45].

Active membrane efflux transport has been shown to be critical to the development of persisters [33, 46-48], the results shown here seem to also point at substrate membrane intake in persisters that occur along logarithmically growing bacteria. One substrate transporter possibly overexpressed in BW25113 but not in bacteria with compromised persistence (*Δfau* and TMP-treated) seems to be *putP*, a proline transporter (Fig. 8). A tempting association of this observation with growth variability based of reported links between the transport of proline via *putP* and *in vivo* growth and pathogenicity in *Staphylococcus aureus* [49-51].

Several studies have used experimental antimetabolites to ameliorate the appearance of bacterial persisters [52-56]. A great advantage of antimetabolites such as antifolates is the fact that they are licensed antimicrobials in clinical practice and have been so for eight decades. Although new antifolates have been added to the cancer chemotherapy pipeline [57], the development and deployment of new antifolates as antimicrobials lags significantly behind [58]. Nonetheless, evidence continues to support the repurposing of antifolates in the treatment of infectious diseases, including those where resistance has become widespread [59-61]. It is encouraging to have found that antifolates affected a known hyper-persister *E. coli* strain as much as the reference strain. This is of relevance for the clinical scenarios where it is likely to find a range of persister levels in active infections. If the appearance of phenotypic variants that persist to antibiotics is closely coupled with the dynamics of exponential generation of cellular mass in *E. coli*, folate metabolism could have a role in the cellular metabolism underlying such variability. Therefore, antifolates could offer a potential avenue for assisting in improving the success of current bactericidal antibiotic treatments.

## Acknowledgements

The project was possible thanks to the support of Liverpool Hope University, School of Health Sciences. Quality control of RNA was implemented with the collaboration of Dr Clare Strode at Edge Hill University, UK.

## Additional Files

Additional file 1 — Inhibitory concentration curves in minimal media of antibiotics and DMSO on the *E. coli* strains used in this study.

Additional file 2 — Inhibitory concentration curves in complex media of antibiotics and DMSO on the *E. coli* strains used in this study.

Additional file 3 — Supplementary tables 1 to 14

Additional file 4 — Supplementary table 15 gene expression BW25113 Additional file 5 — Supplementary table 16 gene expression fau versus BW25113

Additional file 6 — Supplementary table 17 gene expression TMP-treated BW25113 versus BW25113

## Abbreviations

OCFM: folate one-carbon metabolism
5-FCL: 5-formyltetrahydrofolate cyclo-ligase
TMP: trimethoprim
SMX: sulfamethoxazole
DMSO: dimethylsulfoxide
MIQE: Minimum information for publication of quantitative real-time PCR experiments;
GCV: glycine cleavage complex;

